# It’s the fiber, not the fat: Significant effects of dietary challenge on the gut microbiome

**DOI:** 10.1101/716944

**Authors:** Kathleen E. Morrison, Eldin Jašarević, Christopher D. Howard, Tracy L. Bale

## Abstract

**Background:** Dietary effects on the gut microbiome has been shown to play a key role in the pathophysiology of behavioral dysregulation, inflammatory disorders, metabolic syndrome, and obesity. Often overlooked is that experimental diets vary significantly in the proportion and source of dietary fiber. Commonly, treatment comparisons are made between animals that are fed refined diets that lack soluble fiber and animals fed vivarium-provided chow diet that contain a rich source of soluble fiber. Despite the well-established role of soluble fiber on metabolism, immunity, and behavior via the gut microbiome, the extent to which measured outcomes may be driven by differences in dietary fiber is unclear. Further, the significant impact of sex and age in response to dietary challenge is likely important and should also be considered.

**Results:** We compared the impact of transitioning young and aged male and female mice from a chow diet to a refined low soluble fiber diet on body weight and gut microbiota. Then, to determine the contribution of dietary fat, we examined the impact of transitioning a subset of animals from refined low fat to refined high fat diet. Serial tracking of body weights revealed that consumption of low fat or high fat refined diet increased body weight in young and aged adult male mice. Young adult females showed resistance to body weight gain, while high fat diet-fed aged females had significant body weight gain. Transition from a chow diet to low soluble fiber refined diet accounted for most of the variance in community structure and composition across all groups. This dietary transition was characterized by a loss of taxa within the phylum Bacteroidetes and a concurrent bloom of Clostridia and Proteobacteria in a sex- and age-specific manner. Most notably, no changes to gut microbiota community structure and composition were observed between mice consuming either low- or high-fat diet, suggesting that transition to the refined diet that lacks soluble fiber is the primary driver of gut microbiota alterations, with limited additional impact of dietary fat on gut microbiota.

**Conclusion:** Collectively, our results show that the choice of control diet has a significant impact on outcomes and interpretation related to body weight and gut microbiota. These data also have broad implications for rodent studies that draw comparisons between refined high fat diets and chow diets to examine dietary fat effects on metabolic, immune, behavioral, and neurobiological outcomes.

## Background

The increased availability and consumption of energy-dense foods is a key contributor to the global obesity epidemic^1^. The most common rodent model of diet-induced obesity is the consumption of diets that contain 45-60% of energy from dietary fat. This dietary intervention recapitulates key features of metabolic syndrome observed in humans, including body weight gain, increased adiposity, insulin resistance, hyperglycemia, hypertension, and dyslipidemia^2,11–16^. More recently, the gut microbiome has become recognized as a critical intermediary between diet and the metabolic and immune mechanisms contributing to obesity^2–7^. A proposed mechanism linking the gut microbiota to obesity involves high fat diet selection of gut microbial communities with an increased capacity for energy harvest and storage^2,4,5,8,9^. However, the data supporting significant diet and gut microbiota interactions are largely based on comparisons between animals fed diets that are not comparable based on nutritional composition. While high fat diets are formulated as a refined diet in which the source and proportion of every purified ingredient is known and controlled, outcomes in high fat diet-fed animals are most commonly compared to animals fed an unrefined chow diet^2,5,10–12^. Chow diet is a catch-all descriptor for any in-house, institutionally provided vivarium diet. Unlike the refined diets, there is no standardization among chow diets, as nutritional composition is dictated by the market cost of individual ingredients, resulting in variability across batches, lots, and manufacturer^3,4^. Among the various differences between chow and refined diets, the source of dietary fiber is the most important with respect to outcomes on gut microbiome and metabolism^13^. Dietary fibers are broadly classified as soluble or insoluble, with different types in each category. Microbiota ferment soluble fibers to produce short chain fatty acids (SCFAs). In turn, SCFAs provide a major source of energy for colonocytes, promote growth of commensal microbiota and contain outgrowth of pathogenic bacteria, decrease adipose storage, improve insulin sensitivity and decrease local and systemic inflammation^13^. Conversely, insoluble fibers are poorly fermented and therefore do not provide any of the above mentioned benefits^14^. Given that chow diets provide both soluble and insoluble fiber while standard refined diets contain only the insoluble fiber cellulose, the disparity between sources of fiber may have important experimental consequences given their well-established effects on the gut microbiota and metabolism^14–18^.

Moreover, understanding how these dietary components interact with the microbiome has a heightened sense of importance in aging, as the diet of elderly individuals is more likely to be low in fiber^19^. Despite evidence showing that individual differences in response to dietary challenges are driven by age and sex, existing studies have primarily focused on outcomes in young adult males^20–22^. Aging-related alterations in the intestinal microbiota are associated with systemic inflammation in elderly populations, and modulating microbiota community composition and function through dietary intervention has been suggested as a therapeutic method to promote or restore health among aging populations^19,23^. Studies on sex differences in the regulation of metabolism have shown that pre-menopausal women show higher protection against high fat diet-induced obesity relative to men and post-menopausal women^21^. These differences in the susceptibility to obesity and metabolic syndrome are associated with sex differences in circulating gonadal hormones, immunity, and metabolism^19,21,23^. Even in studies that carefully examined age- and sex-specific effects of high fat diet consumption on metabolism and gut microbiota, comparisons were still between mice fed a refined high fat diet to those fed a chow diet^24,25^. Given the compositional differences between unrefined chow and refined diets, it is not clear whether these outcomes are driven by dietary fat or other dietary components such as fiber^15,18,24,25^.

As such, this study sought to examine two overarching hypotheses on the impact of dietary fat and soluble fiber on body weight and gut microbiota. First, to test the hypothesis that differences in soluble fiber drive diet-induced alterations in body weight and gut microbiota, we examined the impact of transitioning young and aged male and female mice from a chow diet to a refined low soluble fiber diet on body weight and gut microbiota. It is important to note that despite the significant differences in dietary fiber composition and content, the refined low fat, low soluble fiber diet (rLFD) and chow diet are comparable based on proportion of dietary fat and caloric density. Second, to test the hypothesis that dietary fat drives changes to body weight and gut microbiota, we examined the impact of transitioning a subset of animals from a 12% fat refined diet (rLFD) to 45% fat, low fiber refined diet (rHFD) on body weight and gut microbiota. As the refined diets are formulated using the same compositionally defined ingredients, the rLFD is a more appropriate control diet for these experiments than the chow diet. We used a serial sampling strategy coupled with 16S rRNA marker gene sequencing to assess longitudinal effects of dietary fiber and fat on gut microbiota community structure and composition. Specifically, we assess the impact transitioning animals from chow to refined diets and the impact of consuming rLFD or rHFD on body weight and gut microbiota. Together, these studies provide insight on critical interactions between diet, sex, and age on the gut microbiota and whole body metabolism.

## Results

### Refined diet promotes weight changes in a sex- and age-specific manner

To test whether progressive removal of soluble fiber is associated with body weight alterations in a sex- and age-specific manner, body weight was tracked in young adult and aged male and female mice across multiple dietary transitions (**Fig. 1A**). All animals were maintained on a chow diet (LabDiet5001) until the initiation of the experiment. Young adult animals consumed the chow diet for 17 weeks and aged animals consumed the chow diet for 60 weeks. As the chow diet is formulated from unrefined ingredients that contains 12% fat and 15% dietary fiber in the form of soluble and insoluble plant polysaccharides, we included a one-week transition period wherein all mice were fed a compositionally defined and refined low fat/low soluble fiber diet (rLFD, Research Diets AIN76A) that contains 12% fat and 5% fiber in the form of insoluble cellulose. Despite the significant differences in dietary fiber composition and content, the rLFD and chow diet are considered to be comparable based on macronutrient profiles and caloric density (**Fig. 1B**)^26^. Following the one week transition period, half of the animals remained on the rLFD while the remaining half was transitioned to a commonly used refined high fat/low soluble fiber diet (rHFD, Research Diets D12451) that contains 45% fat and 5% fiber in the form of insoluble cellulose (**Fig. 1A**). Animals were maintained on rLFD or rHFD for four weeks. As the rLFD and rHFD are formulated using the same compositionally defined ingredients, the rLFD is a more appropriate control diet for these experiments than chow.

**Figure 1.**
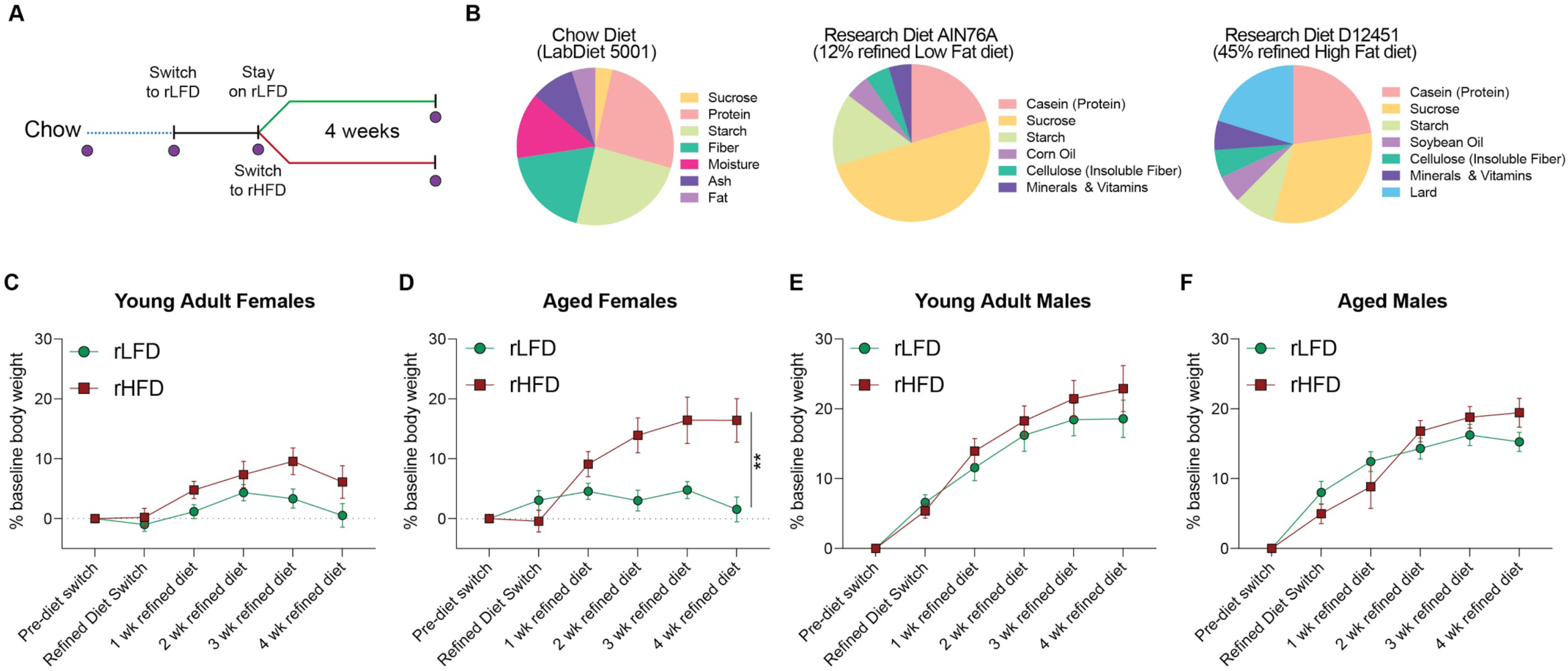
Lack of soluble fiber and increased fat in diet formulations influence weight gain in mice in an age- and sex-specific manner. **(A)** Schematic of experimental study design. Young adult (17 weeks old) and aged (60 weeks old) C57Bl/6:I129 males and females consuming a chow diet were switched to a refined low fat diet (rLFD) for one week to acclimate. Following acclimation to a refined diet, half of the animals remained on rLFD while the other half was switched onto a 45% refined high fat diet (rHFD). Purple circles denote times when fecal samples were collected. Animals were co-housed and therefore all analysis is conducted at the level of the cage to control for co-housing effects (N = 3 cages/age/sex/diet, total N = 92 mice). **(B, C, D)** Composition of diet nutritional composition and ingredients for the **(B)** chow, **(C)** rLFD, and **(D)** rHFD, demonstrating differences in fiber source and quantity between chow and refined diets. **(E-H)** To determine the impact of dietary switching in young adult and aged males and females, weekly body weights were collected prior to refined diet switch, one week following switch to rLFD, and weekly measurements during consumption rLFD or rHFD. **(E)** Body weight was significantly changed over time in young adult females (RM ANOVA, main effect of time, *F*_*5*, 110_ = 15.39, *P* < 0.000, main effect of diet, *F*_*1*, 22_ = 2.920, *P* = 0.1016, N = 24, time*diet interaction, *F*_*5*, 110_ = 2.782, *P* = 0.021). **(F)** Body weight of aged females was significantly changed over time (RM ANOVA, main effect of time, *F*_*5*, 90_ = 17.43, *P* < 0.0001, N = 20), across diets (RM ANOVA, main effect of diet, *F*_*1,18*_ = 6.800, *P* = 0.0178, N = 20), and their interaction (RM ANOVA, time*diet, *F*_*5,90*_ = 12.02, *P* = < 0.0001, N = 20). Post-hoc analysis revealed aged females fed rHFD weighed more at two (*t*_108_ = 3.499, *P* = 0.0041), three (*t*_108_ = 3.748, *P* = 0.0017), and four (*t*_108_ = 4.781, *P* < 0.0001) weeks compared with rLFD-fed aged females. **(G)** Body weight was significantly changed over time in young adult males (RM ANOVA, main effect of time, *F*_*5*, 105_ = 88.146, *P* < 0.0001, main effect of diet, *F*_*1*, 21_ = 0.4240, *P* = 0.522, N = 23). **(H)** Body weight of aged males was significantly changed over time (RM ANOVA, main effect of time, *F*_*5*, 100_ = 67034, *P* < 0.0001, N = 22). Data represented as mean ± SEM. Repeated Measures ANOVA followed by Sidak correction for multiple comparisons. * *P* < 0.05, ** *P* < 0.01, *** *P* < 0.001.

Comparison of weekly body weights across these defined periods of dietary transitions revealed significant age- and sex-specific effects. Comparison between young adult females revealed a main effect of time, no main effect of diet, and a significant interaction between time and diet (RM ANOVA, main effect of time, *F*_*5*,110_ = 15.39, *P* < 0.000, main effect of diet, *F*_*1*,22_ = 2.920, *P* = 0.1016, time*diet interaction, *F*_*5*,110_ = 2.782, *P* = 0.0210, N = 24) Post-hoc analysis revealed that young rHFD females showed a higher percentage of body weight gain at week 5 than young rLFD females, but this effect disappeared by week 6 (Tukey with correction for multiple comparisons; 5 wk, *t*_*20*_ = 2.278, *P* < 0.05) (**Fig 1C**). Body weight comparison in aged females revealed a main effect of time, a main effect of diet and a significant time*diet interaction ((RM ANOVA, main effect of diet, *F*_*1,18*_ = 6.800, *P* = 0.0178, N = 20), and their interaction (RM ANOVA, time*diet, *F*_*5,90*_ = 12.02, *P* = < 0.0001, N = 20). Post-hoc analysis revealed aged females fed rHFD weighed more at two (*t*_108_ = 3.499, *P* = 0.0041), three (*t*_108_ = 3.748, *P* = 0.0017), and four (*t*_108_ = 4.781, *P* < 0.0001) weeks compared with rLFD-fed aged females. These age-specific effects in females may suggest that aged females are more sensitive to lasting high fat diet-induced weight gain (**Fig. 1D**). Males consuming rLFD or rHFD increased body weight across time in a manner that was independent of diet or age (Young adult males, main effect of time, *F*_*5*,105_ = 88.146, *P* < 0.0001, main effect of diet, *F*_*1*,21_ = 0.4240, *P* = 0.522, N = 23; Aged adult males, main effect of time, *F*_*5*, 10_ = 67034, *P* < 0.0001, N = 22; RM ANOVA). These results show that altering sources of fiber, fat, or a combination of both in compositionally defined diets promote weight gain in a sex- and age-specific manner.

### Female-specific changes to fecal microbiota are independent of dietary fat

Alterations to the gut microbiota have been proposed as a key mechanism contributing to high fat diet-induced weight gain and metabolic dysfunction based on studies comparing the gut microbiota of rodents fed a high fat/low soluble fiber diet to those fed a chow diet^2–4,7^. To determine whether the transition from a chow diet that contains soluble fiber to diets that lack soluble fiber and are supplemented with varying proportions of dietary fat influences fecal microbiota, fecal pellets were collected from young adult and aged female mice consuming chow diet, one week following transition to the rLFD and following four weeks of consuming rLFD or rHFD. To examine whether microbiota community structure is altered across dietary transitions, beta and alpha diversity measures were calculated and compared as a function of time consuming rLFD and rHFD diets. Bray-Curtis divergence matrices were calculated to assess the distances between young adult and aged females consuming rLFD and rHFD, and then visualized using PCoA (**Fig. 2A,B**). Permutational multivariate analysis of variance revealed significant effect between chow and refined diets (PERMANOVA, F = 26.284, r^2^ = 0.614, *P* < 0.0001), indicating that one week of consuming a refined diet produced significant community restructuring relative to chow diet females (**Fig. 2A**). Transitioning young adult and aged females from rLFD to rHFD did not further change microbiota community structure (PERMANOVA, F = 1.75, r^2^ = 0.074, *P =* 0.132), indicating that switching female mice from a diet with sufficient levels of soluble fiber (chow) to a diet lacking soluble fiber eclipses the influence of dietary fat on fecal microbiota structure (**Fig. 2B**). Microbial diversity was significantly altered in some (Shannon diversity index) but not the other (observed species counts) indices of alpha diversity in young adult and aged females (Observed species counts, Kruskal-Wallis, H = 9.45, *P* = 0.22; Shannon diversity index, Kruskal-Wallis, H = 26.647, *P* = 0.00038) (**Fig. 2C,D**). The significant changes in the Shannon diversity index was driven by the switch from chow to rLFD, and no additional differences were detected between rLFD and rHFD animals.

**Figure 2.**
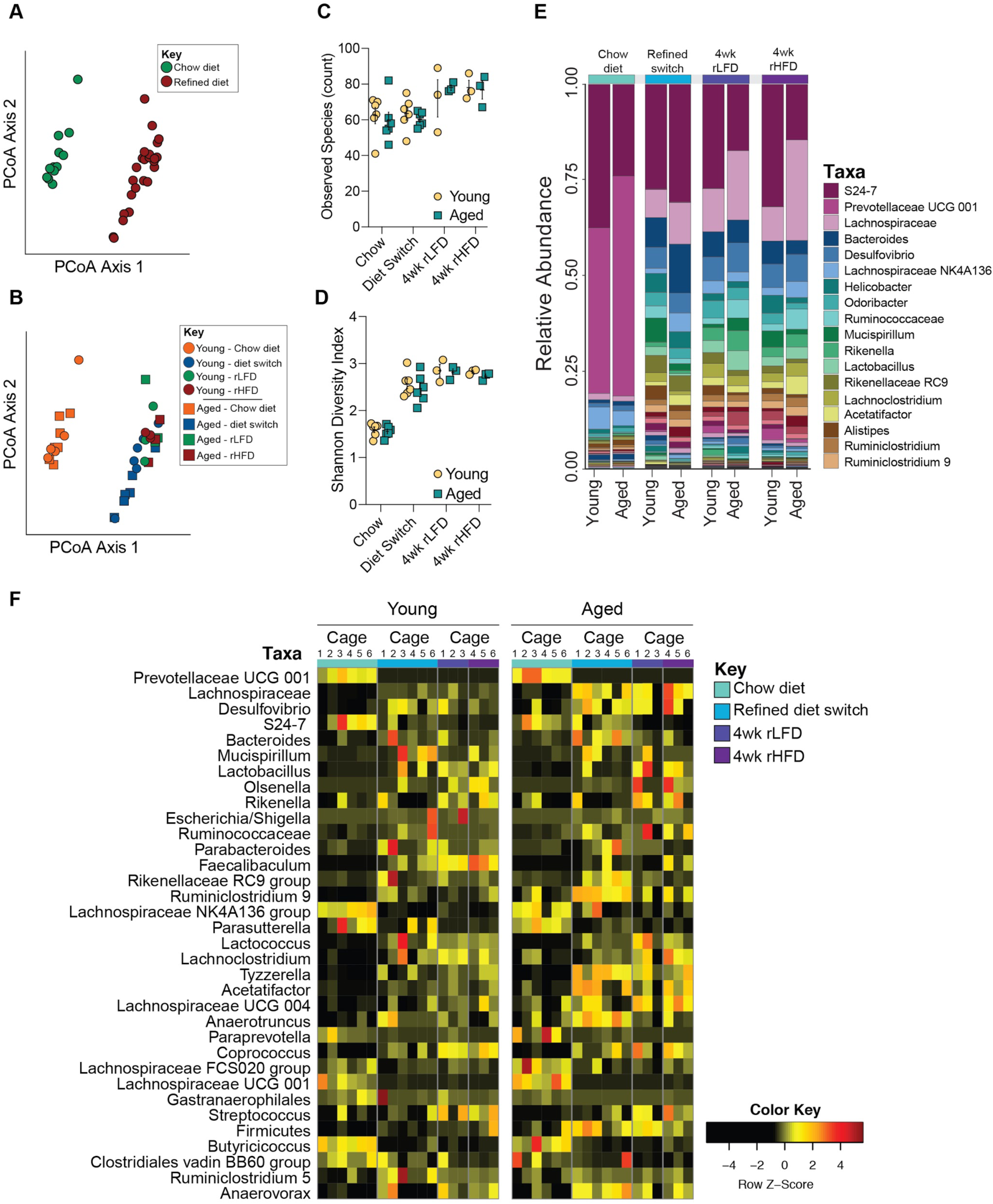
Lack of soluble fiber, not fat, significantly alters gut microbiota composition in young adult and aged female mice. **(A)** Principal coordinates analysis comparing fecal microbiota community structure between females consuming chow and refined diet, demonstrating significant effect of chow and refined diet on community structure (PERMANOVA, F = 26.284, r^2^ = 0.614, *P* < 0.0001), accounting for 67.4% of variance. No difference in community structure between rLFD and rHFD females was observed (PERMANOVA, F = 1.75, r^2^ = 0.074, *P =* 0.132). **(B)** Additional principal coordinate analysis comparing fecal microbiota community structure between young adult and aged females consuming chow, rLFD and rHFD, demonstrating significant interactions of age, rLFD and rHFD on community structure (PERMANOVA, F = 10.122, r^2^ = 0.614, *P* < 0.001). (**C,D**) Comparison of community diversity in young adult and aged females consuming chow, rLFD and rHFD. **(C)** The alpha diversity measure, observed species, plotted against sampling time point demonstrating no impact of diet transition and consumption of refined diets on the number of unique taxa (Kruskal-Wallis, H = 9.45, *P* = 0.22). **(D)** The alpha diversity measure, Shannon diversity index, plotted against sampling time point shows significant differences in community richness and evenness in young adult and aged females (Kruskal-Wallis, H = 26.647, *P* = 0.00038). **(E)** Stacked barplot showing the average relative abundance of taxa in chow, rLFD and rHFD young adult and aged females, characterized by a rapid and lasting loss of taxa within the Bacteroidetes phylum and concomitant bloom of taxa within the Firmicutes and Proteobacteria phyla. Taxa key is truncated to display the 18 most abundant taxa. **(F)** Heatmap depicting 34 significantly different taxa by age and diet in females identified by linear discriminant analysis (FDR < 0.05). Columns represent taxa within each cage. N = 3 female cages/age/diet sampling time point. Data represented as individual data points average per cage ± SEM.

Consistently, comparison of fecal gut microbiota community between chow and refined diets in young adult and aged females revealed significant changes in community composition (**Fig. 2E**). Thus, to determine whether dietary switches impact fecal microbiota composition in young adult and aged female mice, we applied the linear discriminant analysis effect size (LEfSe) method for differential abundance analysis and used a false discovery rate (FDR) cut-off of q < 0.05. LEfSe identified 34 taxa that were affected by switch from chow to refined diets in young adult and aged females (**Fig. 2F**). Comparison of fecal samples from females consuming a chow diet and one week following post-refined diet switch revealed significant decrease of dominant taxa within the phylum Bacteroidetes, including Bacteroidales, Prevotellaceae and S24-7, and taxa within the phylum Firmicutes and order Clostridia, including *Butyricicoccus* (FDR < 0.05) (**Fig. 2F, Supplemental Table 1**). Conversely, transition from the chow diet to rLFD increased abundance of taxa within the phylum Proteobacteria, including *Desulfvibrio* and *Escherichia/Shigella*, and taxa within the phylum Firmicutes and order Clostridia, including *Streptococcus, Mucispirillum, Coprococcus, Olsenella, Ruminoclostridium, Faecalibaculum, Acetatifactor, Rikenella* and *Lactobacillus* (FDR < 0.05). Similarly, switching females from rLFD to rHFD either maintained taxa that were altered during the initial transition from chow to rLFD. Switching females from rLFD to rHFD maintained differences that occurred during the transition from chow to rLFD, suggesting that alterations to gut microbiota composition in young adult and aged females is driven by the lack of soluble fiber in the refined diets rather than proportion of fat (**Fig. 2F, Supplemental Table 1**). In a subsequent analysis, we removed chow diet samples and observed no differences in microbiota structure, diversity or composition between rLFD and rHFD young adult and aged females (all FDR values > 0.05) (**Fig. 2F, Supplemental Table 1**).

Based on our observation that rHFD aged females significantly gain more body weight than aged rLFD females, we next examined whether resistance to weight gain in young adult females and susceptibility to weight gain in aged females is associated with changes to gut microbiota composition. Differential abundance analysis revealed that aged rHFD females had increased abundance of *Bifidobacterium* relative to young adult rHFD females (FDR q = 0.0167). No other differences in gut microbiota composition were detected in females.

### Male-specific changes to fecal microbiota are independent of dietary fat

Similar to the patterns observed in females, community structure of fecal microbiota is significantly different between young adult and aged males following switch from chow to rLFD (PERMANOVA, F = 26.577, r^2^ = 0.617, *P* < 0.0001). No difference in community structure between rLFD and rHFD males was observed (PERMANOVA, F = 1.264, r^2^ = 0.054, *P =* 0.261) (**Fig. 3A,B**). Microbial diversity was significantly different between young adult and aged males (Observed species count, Kruskal-Wallis, H = 17.32, *P* = 0.015; Shannon diversity index, Kruskal-Wallis, H = 26.943, *P* = 0.00034) (**Fig 3C,D**). For both indices of alpha diversity, the significant changes were driven by the switch from chow to rLFD, and no additional differences were detected between rLFD and rHFD animals.

**Figure 3.**
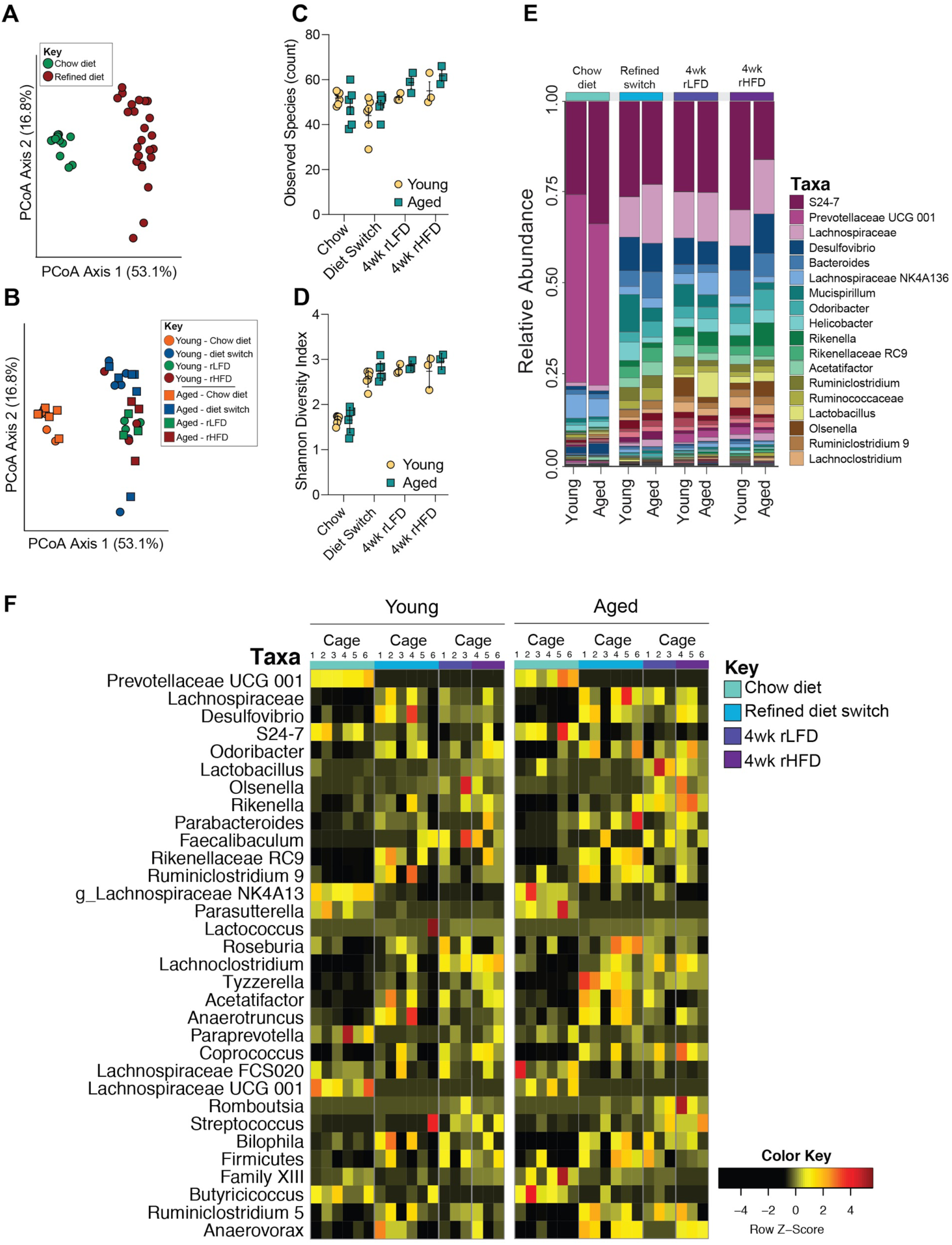
Lack of soluble fiber, not fat, significantly alters gut microbiota composition in young adult and aged male mice. **(A)** Principal coordinates analysis comparing fecal microbiota community structure between males consuming chow and refined diet, demonstrating significant effect of chow and refined diet on community structure (PERMANOVA, F = 26.577, r^2^ = 0.617, *P* < 0.0001), accounting for 69.9% of variance. No difference in community structure between rLFD and rHFD males was observed (PERMANOVA, F = 1.264, r^2^ = 0.054, *P =* 0.261). **(B)** Additional principal coordinate analysis comparing fecal microbiota community structure between young adult and aged males consuming chow, rLFD and rHFD, demonstrating significant interactions of age, rLFD and rHFD on community structure (PERMANOVA, F = 8.84, r^2^ = 0.688, *P* < 0.001). (**C,D**) Comparison of community diversity in young adult and aged males consuming chow, rLFD and rHFD. **(C)** The alpha diversity measure, observed species, plotted against sampling time point demonstrating significant differences in the number of unique taxa following dietary transition and consumption of refined diets (Kruskal-Wallis, H = 17.32, *P* = 0.015). **(D)** The alpha diversity measure, Shannon diversity index, plotted against sampling time point shows significant differences in community richness and evenness in young adult and aged males (Kruskal-Wallis, H = 26.943, *P* = 0.00034). **(E)** Stacked barplot showing the average relative abundance of taxa in chow, rLFD and rHFD young adult and aged males, characterized by a rapid and lasting loss of taxa within the Bacteroidetes phylum and concurrent bloom of taxa within the Firmicutes and Proteobacteria phyla. Taxa key is truncated at the 18 most abundant taxa. **(F)** Heatmap depicting 32 significantly different taxa by age and diet in females identified by linear discriminant analysis (FDR < 0.05). Columns represent taxa within each cage. Data represented as individual data points averaged per cage ± SEM.

Consistent with alterations to community structure and diversity, the fecal microbiota between chow and refined diets revealed significant remodeling (**Fig. 2E**). Differential abundance analysis by LEfSe identified 32 taxa that differed between chow and refined diet-fed males. Transition from chow to rLFD resulted in a significant decrease in taxa within the Bacteroidetes phylum, including Prevotellaceae and Lachnospiraceae, and taxa within the Firmicutes phylum, including *Butyricicoccus* (FDR < 0.05) (**Fig. 3E, Supplemental Table 2**). Concurrently, we observed a significant increase in the abundance of taxa within the Proteobacteria phylum, including Desulfvibrio and Parasutterella, and taxa within the phylum Firmicutes, including *Roseburia, Coprococcus, Bilophila, Olsenella, Faecalibaculum, Acetatifactor, Rikenella* and *Lactobacillus* (FDR < 0.05). Similar to females, switching males from rLFD to rHFD maintained differences in taxa that were altered during the switch from chow to rLFD. In a subsequent analysis, we removed chow diet samples and observed no differences in microbiota structure, diversity or composition between rLFD and rHFD young adult and aged males (all FDR values > 0.05) (**Fig. 2E, Supplemental Table 2**). No other differences in gut microbiota composition were detected in males. Taken together, these results demonstrate that removal of soluble fiber is an important driver of gut microbiota alterations, while varying proportions of dietary fat may exacerbate these changes.

### Sex-specific effects on gut microbiota composition following transition to refined diets

Given the significant interaction between sex, age, and diet on gut microbiota community structure and composition, we next examined whether the gut microbiota composition between males and females differed following dietary transitions. Additional analysis revealed that males and females shared 27 taxa that were differentially abundant following transition to refined diet (FDR < 0.05) (**Supplemental Figure 2A**). Seven taxa were differentially abundant in young adult and aged females, while five taxa were differentially abundant in young adult and aged males (FDR < 0.05) (**Supplemental Figure 2B-G**). These results contribute to a growing literature demonstrating sex-specific diet effects on gut microbiota community and composition and highlight important sex and age interactions on diet, gut microbiota and body weight.

## Discussion

Dietary intervention studies largely revolve around altering fat content, as diet-induced obesity is comorbid with a variety of serious health conditions in humans, including metabolic disorders, type 2 diabetes, heart disease, liver disease, mood disorders, and cancer^31–33^. However, rodent and nonhuman primate studies have overlooked other components of diet that could contribute to disease risk, most notably fiber^34^. In the “Westernized diet” that is often the target of these studies, little consideration has been given to the amount of fiber and whether or not it is soluble. Studies that do account for differences in fiber source in diet formulations demonstrate that high fat diet-induced obesity in young adult male mice is driven by the lack of soluble fiber in refined diet formulations^16,34,35^. Furthermore, other potentially important dietary components have not been considered, therefore limiting the capacity to isolate the effects of the dietary challenge specifically to the fat content. This is a major bottleneck in understanding how diet is involved in the pathophysiology of metabolic and related disorders. The literature on age- and sex-specific effects of high fat diet consumption on metabolism and gut microbiota is subsequently based on studies comparing mice fed a high fat/low soluble fiber diet to those fed a chow diet^36–38^. Thus, we examined the age- and sex-specific effect of a refined high fat/ow fiber diet (rHFD) on body weight and gut microbiota composition relative to mice fed a refined low-fat diet (rLFD) that is nutritionally and compositionally matched to the rHFD.

To determine the contribution of soluble fiber and dietary fat on whole body metabolism, body weight measurements were collected from young adult and aged male and female mice while consuming chow and refined diets. After 4 weeks, young adult female mice showed resistance to weight gain to rHFD, consistent with previous reports^29,39,40^. Conversely, aged females fed rHFD showed rapid body weight gain relative to rLFD-fed aged females. Age-specific vulnerability to diet-induced body weight gain in females may be related to aging related changes to estrogens. Circulating estrogens have been shown to be protective against diet-induced obesity by promoting glucose tolerance and insulin resistance, as well as engaging in the direct inhibition of proinflammatory cytokines that are stimulated by consumption of a high fat diet^39–41^. Reduction in circulating estrogens through ovariectomy dramatically increases risk for high fat diet-induced obesity and metabolic disorders in mice and post-menopausal women, providing a possible explanation for rHFD-induced weight gain in aged females^21^.

Conversely, young adult and aged males showed a significant gain in body weight that was independent of refined diet formulation. The significant increase in body weight in young adult and aged males fed either rLFD or rHFD may suggest that other components of the refined diet contribute to body weight gain that is independent of dietary fat^16,17,34^. Consistent with our observations, increased body weight and adiposity, loss of intestinal mass and decreased availability of short chain fatty acids has been previously reported in young adult male mice fed the rLFD relative to chow diet-fed males^16,34^. Taken together, these results suggest that a lack of soluble fiber and varying proportions of dietary fat exhibit significant effects on body weight in an age- and sex-specific manner, and further highlights the importance of an appropriate refined diet control for diet-induced obesity studies assessing metabolic outcomes.

High fat diet-induced weight gain and metabolic dysfunction have been mechanistically linked to altered gut microbiota composition and functional capacity to harvest energy, however, these conclusions are drawn from studies comparing the gut microbiota of rodents fed a high fat/low soluble fiber diet to those fed a chow diet ^2,5,6,6,11,41^. To determine whether switching mice from a chow diet that contains soluble fiber to low fiber, varied fat content refined diets influences fecal microbiota, fecal pellets were collected from young adult and aged male and female mice while consuming chow diet, one week following transition to the rLFD and following four weeks of consuming rLFD or rHFD. Transition from the chow diet to rLFD resulted in changes to microbiota community structure and composition in all groups, regardless of sex and age. This dietary transition was characterized by a loss of taxa within the phylum Bacteroidetes and a concomitant bloom of Clostridia and Proteobacteria in a sex- and age-specifc manner. Specifically, the bloom of *Escherichia/Shigella* in females, but not males, is an interesting observation for future investigation given the interaction between poor nutritional environment of the gut, bloom of Proteobacteria, and systemic inflammation^42^. Similar changes to gut microbiota have been previously reported in mice consuming a rHFD, suggesting that the bloom of Clostridia and Proteobacteria may be attributed to loss of soluble fiber rather than the high proportion of dietary fat^41^. Further, dramatic changes in gut microbiota composition within one week of switching from a chow diet to the rLFD highlight the exquisite sensitivity of the gastrointestinal ecosystem to shifts in nutrient availability and composition^43,44^. Consistent with recent reports, the lack of differences between rLFD and rHFD-fed mice may indicate that gut microbiota structure and composition can be dissociated from body weight and systemic inflammation^17,34,35^. Further, these results suggest that the absence of soluble fiber in both refined diets exerts a more dominant effect than the varying proportions of fat and carbohydrates between these diets.

Collectively, our results highlight that the choice of control diet for diet-induced metabolic disease experiments may influence interpretations related to gut microbiota composition. These data also have broad implications for rodent studies that draw comparisons between refined high fat/low soluble fiber diets and chow diets to examine dietary fat effects on metabolic, immune, behavioral, and neurobiological outcomes. Moving forward, it will be of great interest to examine the mechanisms underlying the sex differences of soluble fiber in whole body metabolism and determine whether compositionally defined combinations of fiber and fat influence metabolism, immunity, and neural circuits that control feeding behavior and energy homeostasis.

## Methods

### Ethical Approval and Institutional Governance

All experiments were approved by the University of Pennsylvania Institutional Animal Care and Use Committee and performed in accordance with National Institutes of Health Animal Care and Use Guidelines.

### Animals

Male and female mice used in this study were derived from an in-house C57Bl/6:129 mixed strain. Mice were housed in a 12 hour light:dark cycle. Food (LabDiet 5001) and water were provided *ad libitum*. Mice were group housed following weaning at postnatal day 28.

### Experimental Design

Figure 1A provides an overview of the experimental design. All mice were maintained on Chow Diet (LabDiet 5001) and fresh fecal pellets were collected prior to refined diet transition. Two refined diets were used during the experiment: refined low fat/low soluble fiber diet (rLFD, Research Diets AIN76A) supplying energy as 12% fat, 21% protein, and 67% carbohydrates; and refined high fat/low soluble fiber diet (rHFD, Research Diets D12451) supplying energy as 45% fat, 20% protein, and 35% carbohydrates. First, all mice were placed on the rLFD for one week to acclimate them to a refined diet. Following acclimation, mice were randomly to either remain on rLFD or switch to rHFD. At that point, a total of 12 young adult males, 12 young adult females, 11 aged males and 10 aged females remained on rLFD and a total of 12 young adult males, 12 young adult females, 12 aged males and 11 aged females were placed on rHFD. Diets were available for *ad libitum* consumption. Body weight was recorded once per week. Fecal pellets were collected following the 1 week acclimation period and 4 weeks post-diet transition. While individual fecal pellets were collected, mice remained group housed for the entire course of the experiment. As group housing condition has been shown to account for a significant portion of variance in fecal microbiota community composition, all analysis was conducted at the level of the cage (N = 3 cages per sex/diet).

### DNA extraction

Genomic DNA from fecal samples were isolated using the Stratec PSP Spin Stool DNA Plus kit using the difficult to lyse bacteria protocol from the manufacturer (STRATEC Molecular GmbH, Berlin, Germany). Each sample DNA was eluted into 100 µL of Elution Buffer provided by the Stratec PSP Spin Stool DNA Plus kit.

### Illumina MiSeq 16S rRNA marker gene sequence data processing and analysis

The V4 region of the bacterial 16S rRNA gene was amplified using a dual-index paired-end sequencing strategy for the Illumina platform as previously described^45^. Sequencing was performed on a MiSeq instrument (Illumina, San Diego, CA) using 2×250 base paired-end chemistry at the University of Maryland School of Medicine Institute for Genome Sciences. The sequencing run yielded a total of 25,260,257 read counts with an average of 33,292 read counts per sample. The sequences were demultiplexed using the dual-barcode strategy, a mapping file linking barcode to samples and split_libraries.py, a QIIME-dependent script^46^. The resulting forward and reverse fastq files were split by sample using the QIIME-dependent script split_sequence_file_on_sample_ids.py, and primer sequences were removed using TagCleaner (version 0.16)^47^. Further processing followed the DADA2 workflow for Big Data and DADA2 (v.1.5.2) (https://benjjneb.github.io/dada2/bigdata.html)^48^. Data filtering was set to include features where 20% of its values contain a minimum of 4 counts. In addition, features that exhibit low variance across treatment conditions are unlikely to be associated with treatment conditions, and therefore variance was measured by inter-quartile range and removed at 10%. Data was normalized by cumulative sum scaling and differential abundance analysis was conducted using Linear Discriminant Analysis effect size with an FDR cut-off at q < 0.05^49^. For quality control purposes, water and processed blank samples were sequenced and analyzed through the bioinformatics pipeline. Taxa identified as cyanobacteria or ‘unclassified’ to the phylum level were removed. The sequencing data has been deposited in the Sequence Read Archive (SRA) of the National Center for Biotechnology Information (NCBI) (Bioproject: PRJNA6083825) to be released upon publication.

### Statistical analysis

Data are represented as individual data points or individual points averaged per cage ± SEM. Body weight data were analyzed by repeated measures analysis of variance and post-hoc analysis was conducted using Sidak correction for multiple comparisons. Given the nonparametric nature of microbiota data, indices of alpha diversity data was analyzed using Kruskal-Wallis test. Permutational multivariate analysis of variance was used to analyze effects of diet, sex and age. 16S rRNA marker gene sequencing raw count data was filtered was set to include features where 20% of its values contain a minimum of 4 counts. In addition, features that exhibit low variance across treatment conditions are unlikely to be associated with treatment conditions, and therefore variance was measured by inter-quartile range and removed at 10%. Data was normalized by cumulative sum scaling and differential abundance analysis was conducted using Linear Discriminant Analysis effect size with an FDR cut-off at q < 0.05. Analysis was conducted in the R environment and Bionconductor using the metagenomeSeq package, heatmaps were visualized using the heatmap.2 function within the gplots package, and bar plots were visualized using GraphPad^50–53^.

## Acknowledgments

We thank Jacques Ravel, Mike Humphrys and Luke Tallon from the Institute for Genome Sciences at the University of Maryland School of Medicine for assistance with 16S rRNA marker gene sequencing. T.L.B was supported by the National Institutes of Health under Award Numbers P50-MH099910, MH 104184, MH 091258, MH 087597, MH 073030, MH 108286, HD 097093, and ES 028202. E.J. was supported by the National Institutes of Health National Research Service Award F32 MH 109298. K.E.M. was supported by National Institutes of Child Health and Human Development Pathway to Independence Award K99 HD 091376.

---

**Supplemental Figure 1.**
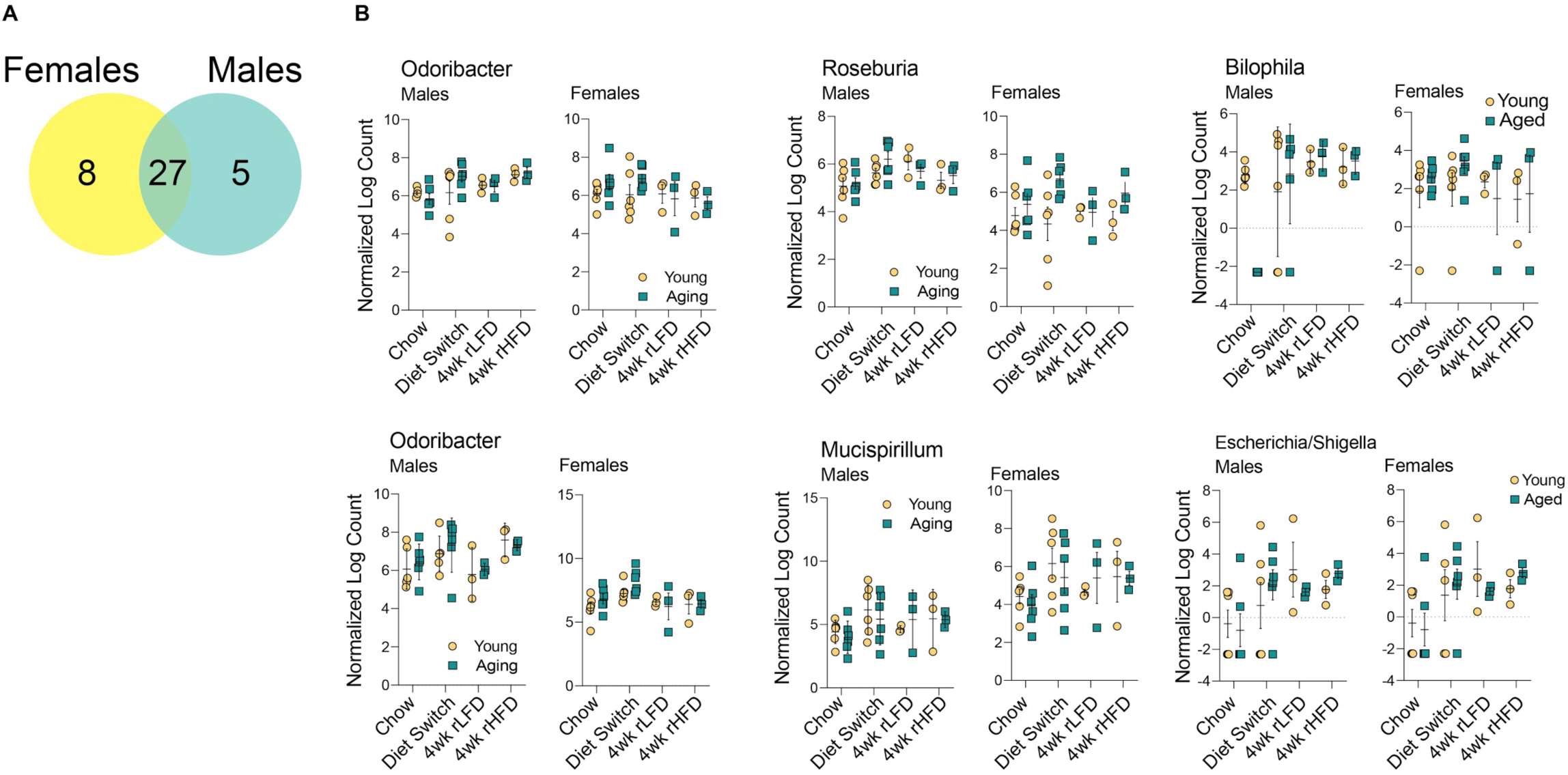
Sex-specific alterations of refined diet on gut microbiota composition. **(A)** Venn diagram depicting microbiota significantly altered by diet in young adult and aged males and females (27), female-specific taxa altered by diet (7) and male-specific taxa altered by diet (5) identified by linear discriminant analysis (FDR < 0.05, N = 3 cages/sex/age/diet sampling timepoint). **(B-G)** Taxa abundance plotted against sampling time point showing sex-specific differences identified by linear discriminant analysis (FDR = 0.05). Data represented as individual data points averaged per cage ± SEM.

**Supplemental Table 1. Results from linear discriminant analysis of microbiota composition in female mice**.

**Supplemental Table 2. Results from linear discriminant analysis of microbiota composition in male mice**.

